# Treatment of human cells with 5-aza-dC induces formation of PARP1-DNA covalent adducts at genomic regions targeted by DNMT1

**DOI:** 10.1101/538645

**Authors:** Kostantin Kiianitsa, Nancy Maizels

## Abstract

The nucleoside analog 5-aza-2’-deoxycytidine (5-aza-dC) is used to treat some hematopoietic malignancies. The mechanism of cell killing depends upon DNMT1, but is otherwise not clearly defined. Here we show that PARP1 forms covalent DNA adducts in human lymphoblast or fibroblasts treated with 5-aza-dC. Some adducts recovered from 5-aza-dC-treated cells have undergone cleavage by apoptotic caspases 3/7. Mapping of PARP1-DNA adducts, by a new method, “Adduct-Seq”, demonstrates adduct enrichment at CpG-dense genomic locations that are targets of maintenance methylation by DNMT1. Covalent protein-DNA adducts can arrest replication and induce apoptosis, and these results raise the possibility that induction of PARP1-DNA adducts may contribute to cell killing in response to treatment with 5-aza-dC.

## 1. Introduction

The nucleoside analog 5-aza-2’-deoxycytidine (5-aza-dC; clinically known as decitabine) is incorporated opposite G during DNA replication. Incorporation of 5-aza-dC opposite 5-methyl-CpG creates a target for methylation by the maintenance DNA methyltransferase, DNMT1. Treatment of mammalian cells with 5-aza-dC also causes proteolysis of DNMT1 that results in genomewide loss of cytosine methylation and extensive epigenetic reprogramming [1-3]. This gave rise to the notion that 5-aza-dC might re-activate silenced tumor suppressor genes to limit cell proliferation, and 5-aza-dC is used in the clinic as an “epigenetic drug” to treat myelodysplasia syndrome (MDS) and acute myelogenous leukemia (AML) in patients who are unlikely to tolerate other treatments [4]. However efforts to demonstrate that 5-aza-dC efficacy correlates with reprogramming of specific genetic networks or pathways have not been fruitful [5]. This prompted us to further investigate the possibility that cell killing results from DNA damage induced by 5-aza-dC treatment.

DNMT1 normally forms an obligatory but transient covalent bond with its target in the course of transfer of a methyl group from S-adenosyl methionine to DNA [6]. At a 5-aza-substituted site, the DNMT1-DNA adduct persists, resulting in base damage, genomic instability or cell killing; and deficiency in DNMT1 provides protection from these negative outcomes of 5-aza-dC treatment [7-12]. PARP1 is a conserved and abundant factor involved in repair and transcriptional regulation which is recruited to many kinds of damage, including base damage like that resulting from 5-aza-dC treatment [12-16]. PARP1 has been shown to form covalent adducts with DNA substrates containing AP sites or 5’-dRP residues in vitro, and in living cells treated with an alkylating agent [17-19]. Persistent covalent protein-DNA adducts can cause replication arrest and induce apoptosis. This led us to hypothesize that recruitment of PARP1 to damage resulting from aberrant DNMT1 activity at 5-aza-substituted cytosines might cause PARP1-DNA adducts to form and then persist, arresting replication and inducing apoptosis.

To test this, we have isolated PARP1-DNA adducts from human cells treated with 5-aza-dC and characterized adducted PARP1 by Western blotting and adducted DNA by genomic sequencing. Here we report that PARP1-DNA adducts can be readily recovered from 5-aza-dC treated cells. Caspase cleavage of PARP1 is a hallmark of apoptosis [20-22], and some PARP1 adducts carried the signature N-terminal neo-epitope generated upon caspase cleavage. Adducted DNA fragments were sequenced using a new method, “Adduct-Seq”, which demonstrated highly significant enrichment at genomic features characteristic of the CpG islands (CGIs) that are targets of methylation by DNMT1. Caspase-cleaved PARP1 is impaired for recruitment to chromatin [23], and while we cannot rule out the possibility that caspase cleavage of PARP1 occurred prior to adduct formation, our results better support a model in which aberrant activity of DNMT1 at 5-aza-substituted regions creates damage that recruits PARP1 to form covalent adducts, which arrest replication and kill cells.

## 2. Materials and Methods

### 2.1 Cells, cell culture, and drug treatment

K562 lymphoblasts (ATCC CCL-243) were obtained from ATCC and cultured in RPMI medium with 10% fetal calf serum. GM639 cells (gift of Dr. Ray Monnat, University of Washington), were cultured in DMEM medium with 10% fetal calf serum. 5-aza-dC was obtained from Calbiochem/EMD Millipore; the PARP inhibitor BMN-673 (talazoparib), from ApexBio.

### 2.2 Antibodies

Antibodies used for Western blots included: anti-rec-PARP1, a rabbit polyclonal raised against recombinant PARP1 (Enzo Life Sciences ALX-210-302-R100; 1:4000 dilution); anti-N-ter-PARP1, a rabbit polyclonal raised against the N-terminal half of recombinant PARP1 (Active Motif Ab 2793257, 1:2000 dilution); anti-C-ter-PARP1, an affinity-purified mouse mAb that recognizes an epitope in the C-terminal NAD binding and catalytic domain of human PARP1 (clone 7D3-6, BD Biosciences #556493; 1:500 dilution); anti-cc-PARP1, a mouse mAb specific for the neo-epitope generated at the N-terminal of the 89 kD PARP1 fragment formed after cleavage by apoptotic caspases between Asp214/Gly215 (clone F21-852, BD Biosciences #552596, 1:1000 dilution). Secondary detection was with HRP-conjugated goat anti-mouse IgG or donkey anti-rabbit IgG (BioLegend, #405306 and #406401, respectively; 1:5000 dilution). Immune complexes were visualized using the SuperSignal West system (Pierce). Specificity of PARP1 Abs for full-length 113 kD PARP1 and its cleavage products was assessed by Western blots of whole cell extracts treated with staurosporine (EMD Millipore; 1 µM, 1 hr), a kinase inhibitor that induces apoptotic PARP1 cleavage (**Supplementary Fig. S1B**).

### 2.3 Nucleic acid and protein quantification

DNA and RNA were quantified using DNA/RNA specific fluorescent detection kits (Qubit, Invitrogen). Total protein was measured using BCA detection kit (Pierce).

### 2.4 RADAR/Western analysis

DNA-protein adducts were purified for RADAR/Western analysis using an adaptation of the RADAR (<u>R</u>apid <u>A</u>pproach to <u>D</u>NA <u>A</u>dduct <u>R</u>ecovery) fractionation protocol in which cells or nuclei are directly lysed in the presence of chaotrope and detergents, after which nucleic acids including protein-DNA adducts are separated from total protein by precipitation in alcohol [24, 25]. Preparations used 2-3 x 10^7^ K562 cells or 1.5-2.5 x 10^7^ GM639 cells. Cells were harvested by centrifugation and washed in PBS. Cytoplasm, including rRNA and up to 75% of total protein was eliminated by quick lysis in 1 ml MPER reagent (Pierce/Thermo Fisher) supplemented with Halt protease inhibitor cocktail (Thermo Fisher) and 1 mM DTT. Homogenization was performed by 8-10 pipet strokes, then nuclear pellets collected by 2 min centrifugation at maximal speed. PARP1 and DNMT1 were confirmed to be exclusively in the pellet by Western blotting; as was essentially all genomic DNA. Pellets were promptly lysed in 400 µl of RADAR lysis buffer (LS1), consisting of 5 M guanidinium isothiocyanate (GTC), 2% Sarkosyl, 10 mg/ml DTT, 20 mM EDTA, 20 mM Tris-HCl (pH 8.0) and 0.1 M sodium acetate (pH 5.3), and adjusted to final pH 6.5 with NaOH. Lysates were transferred to polystyrene tubes (Evergreen) and sonicated on ice using a cup horn device (QSONICA) with ten 30 sec pulses at amplitude 100, and 60 sec cool-off time between pulses. Sonication yielded an average DNA size of 500 bp. After sonication, lysates were mixed with 80 µl 12 M LiCl (final concentration 2M LiCl, 4.16 M GTC) and incubated for 10 min at 37°C on a Thermoshaker, then cleared by 10 min centrifugation at 14,000 rpm at room temperature prior to conservative recovery of the supernatant. Lysates (volume approximately 450 µl) were transferred to new tubes, mixed with an equal volume of isopropanol; and nucleic acids were precipitated by 5 min centrifugation at 14,000 rpm at room temperature. Pellets were washed thrice with 1 ml 75% ethanol, followed by 2 min centrifugation. Nucleic acid pellets were dissolved in 100-200 µl of freshly prepared 8 mM NaOH by 30-60 min shaking on a Thermomixer at 37°C; then neutralized by addition of 1 M HEPES to a final concentration of 20 mM. The resulting samples contained mostly genomic DNA and some nuclear RNA, with a yield of approximately 70 µg DNA per 10^7^ cells. The DNA/protein weight ratio was 100:3 or higher, comparable to adducts prepared by ultracentrifugation in a CsCl gradient [26].

Prior to gel electrophoresis, samples containing 10-30 µg genomic DNA were treated with Benzonase nuclease (EMD Millipore; 0.5 units per µg DNA) in the presence of 2 mM magnesium chloride for 30 min at 37°C to digest DNA. Completeness of nuclease digestion was verified by Qubit assays. Samples were resolved on precast 4-12% gradient protein gels (Bolt, Invitrogen), transferred to nitrocellulose membranes (0.45 µM pore, Invitrogen) and probed with antibodies in PBST buffer (PBS containing 0.05% Tween 20) with 0.5% alkali-soluble casein (Novagen) as blocking agent.

### 2.5 Adduct-Seq analysis

To identify genomic locations of adducted PARP1, we developed an approach, “Adduct-Seq” which resembles ChIP-Seq but depends upon adduction rather than formaldehyde-induced crosslinking to form stable protein-DNA complexes (**Fig. 2A**, right). An analogous approach has previously been used to enrich genomic DNA adducted to Topoisomerase 1 [27]. In brief, adducts were first isolated from approximately 1.5 x 10^7^ untreated or treated cells by RADAR fractionation, sonicated as above to reduce the size of adducted DNA; PARP1-DNA adducts captured by anti-PARP1 mAbs bound to a solid support; and adducted DNA fragments released by Proteinase K treatment and sequenced. To avoid melting of duplex DNA that may later interfere with adaptor ligation for Illumina sequencing, DNA was resuspended with 5 mM tris-HCl (pH 8.5) rather than 8 N NaOH. Antibody capture was performed using a ChromaFlash High-Sensitivity ChIP kit (Epigentek) according to the manufacturer’s instructions. Briefly, strip wells were pre-coated with anti-C-ter-PARP1 mAb (see above; 800 ng IgG per well), unbound mAbs removed by washing, then incubated with 40 µg RADAR-fractionated DNA overnight at 4°C in 100 µl of provided buffer, washed, treated with RNase A and rewashed. DNA was released by treatment with Proteinase K and purified using a ChIP DNA Clean & Concentrator kit (Zymo Research). DNA yield was approximately 1-2 ng/10^7^ cells, and 1.5 ng DNA was subject to paired end Illumina sequencing, 100 nt per read, with ∼40 million read pairs per sample sequencing depths.

**Fig. 1.**
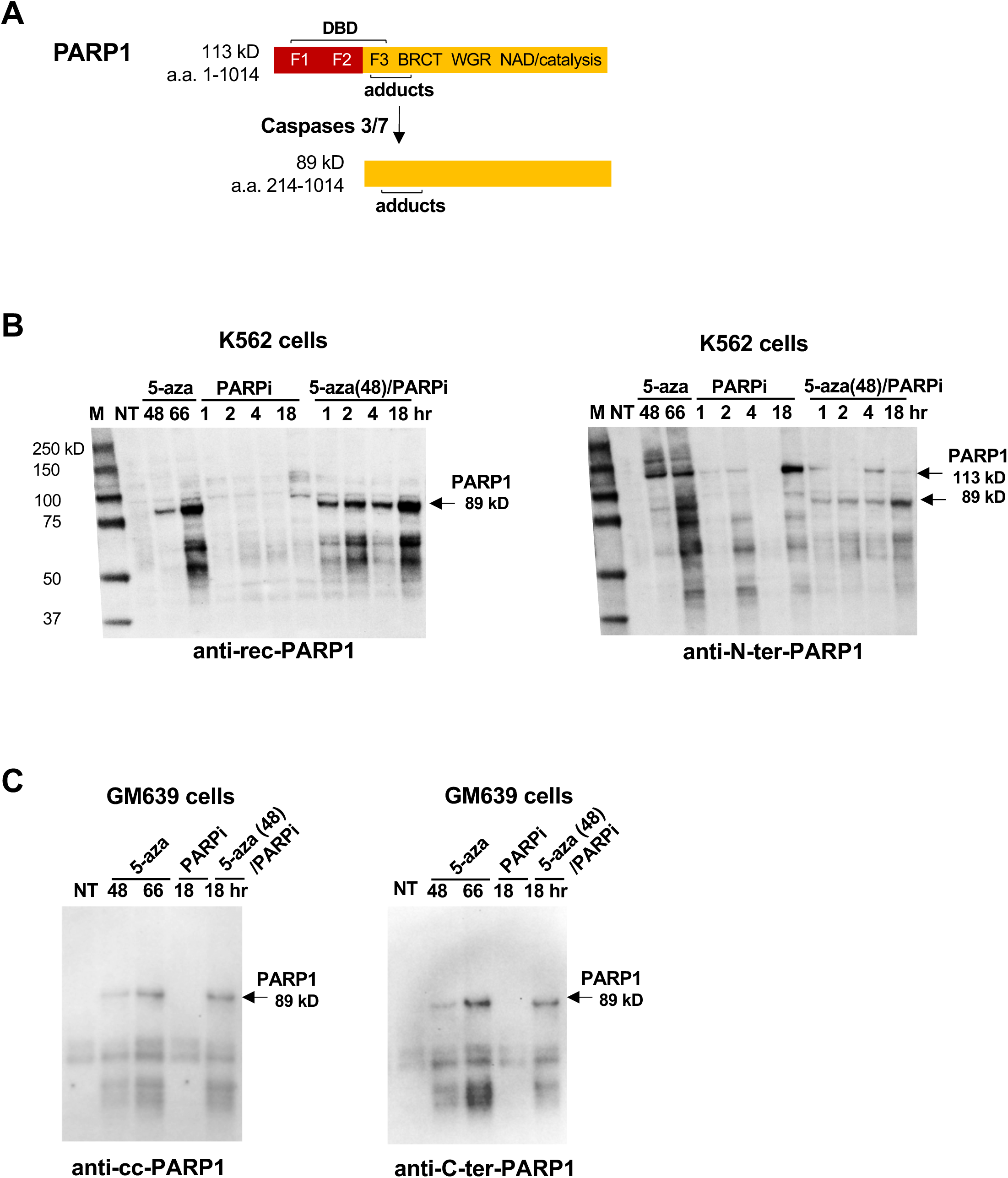
RADAR/Western blots Identify PARP1-DNA Adducts Induced by 5-aza-dC. (A) PARP1 apoptotic cleavage. The 113 kD PARP1 polypeptide contains three zinc finger motifs (F1, F2 and F3) in the DNA binding domain (DBD); the BRCT, WGR and NAD binding/catalytic domains; and the region to which covalent adducts have been mapped [18]. Caspases 3/7 cleave PARP1 within the DBD, generating a 24 kD fragment (not shown) and an 89 KD fragment bearing a new N-terminal neo-epitope. (B) Western blot analysis of PARP1-DNA adducts isolated from K562 cells, probed with (left) anti-rec-PARP1 or (right) anti-N-ter-PARP1. A single blot was washed but not stripped between probings. Cells were treated for indicated times with 1 µM 5-aza-dC (5-aza) or1 µM BMN-673 (PARPi); or pretreated with 1 µM 5-aza-dC for 48 hr, after which PARPi was added and culture continued for indicated times. M, marker (kD); NT, untreated sample. Arrows identify 113 kD and 89 kD PA. (C) Western blot analysis of PARP1-DNA adducts isolated from GM639 cells. Duplicate blots were probed with (left) anti-cc-PARP1 mAb or (right) anti-C-ter-PARP1 mAb. Notations as in panel B.

**Fig. 2.**
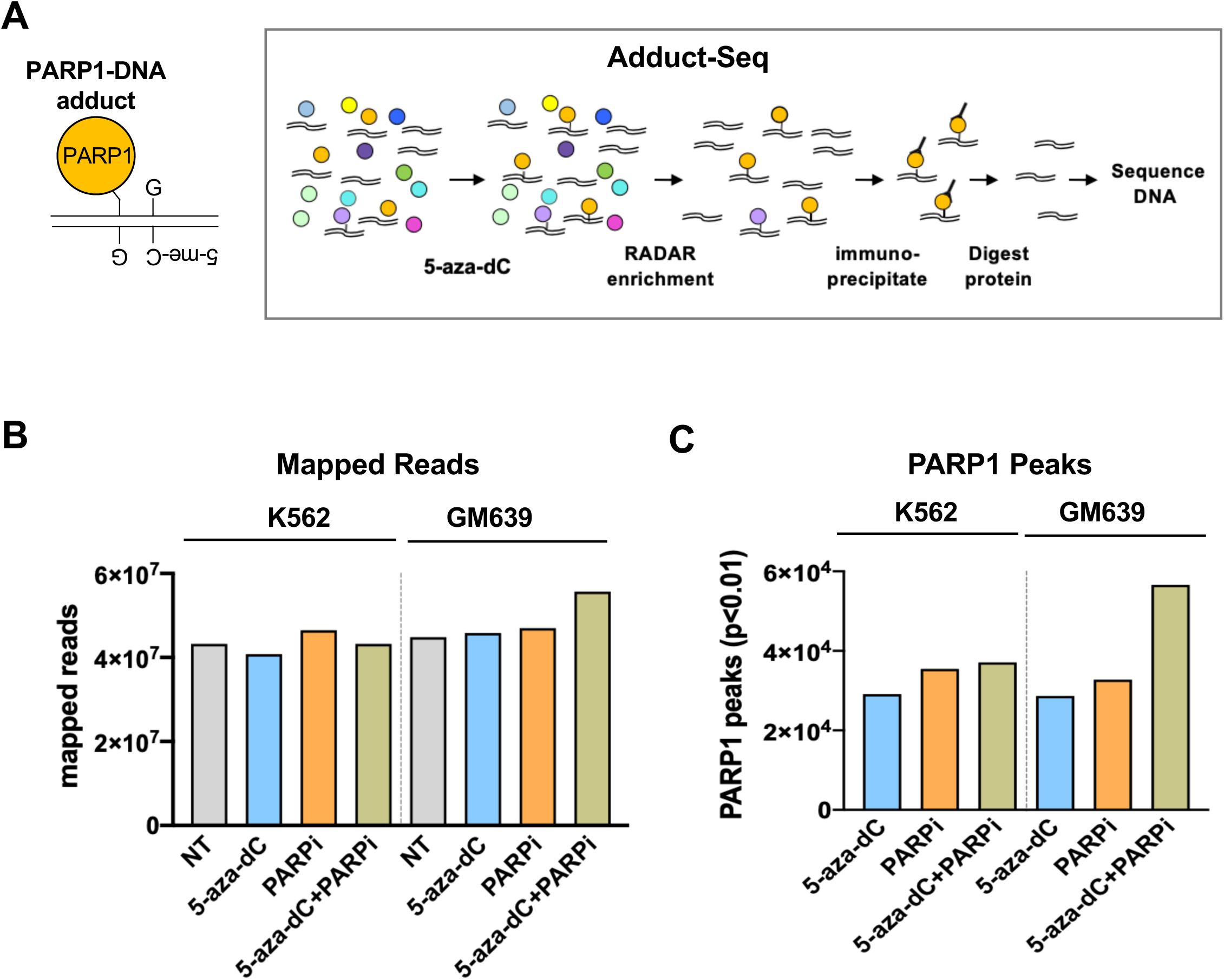
Recovery of PARP1-DNA adducts by Adduct-Seq. (A) Outline of Adduct-Seq characterization of adducted DNA. Left, Predicted PARP1-DNA adduct at hemi-methylated CpG. Right, recovery of PARP1-DNA adducts; see text for details. Protein (circles), DNA (double lines). (B) Mapped reads generated by Adduct-Seq of K562 or GM639 cells untreated (NT) or treated as indicated. (C) PARP1 peaks identified by Adduct-Seq in K562 or GM639 cells treated as indicated.

Raw reads were mapped to human genome (hg38) using Bowtie2. Genomic locations (peaks) of adducted PARP1 in response to treatment were identified by MACS2 from mapped reads of Adduct-Seq libraries (“narrow peak” calling option, p<0.01 cutoff) using the untreated sample to assess background. The GenometriCorr R package (https://doi.org/10.1371/journal.pcbi.1002529 [28]) was used to test spatial correlation of PARP1 peaks and 31,144 CGIs available as standard track on the UCSC Genome Browser. Multiple tests were performed to assess spatial correlation: absolute and relative distance between query (PARP1 peaks) and reference (CGIs), Jaccard test (union vs intersection for each reference feature) and projection test (overlap of query with reference features). GC fraction and CpG density of query and reference sequences were calculated using R functions gcContent and cpgDensity, respectively.

### 2.6 Statistical analyses

Graphing, calculation of mean values and statistical tests of significance were performed using GraphPad Prism software.

## 3.Results

### 3.1 RADAR/Western blots identify PARP1-DNA adducts induced by 5-aza-dC

The possibility that PARP1 might contribute to cell killing made it important to not only ask if PARP1-DNA adducts formed in cells treated with 5-aza-dC, but to characterize those adducts. The 113 kD PARP1 polypeptide (**Fig. 1A**) contains an N-terminal DNA binding domain (DBD) and a C-terminal domain that catalyzes polymerization of poly(ADP-ribose) (PAR) from the coenzyme, NAD^+^, enabling PARP1 to modify itself and other proteins by addition of chains of PAR (“PARylation”) which stimulates protein recruitment for DNA repair. In apoptotic cells, activated caspases 3/7 cleave 113 kD PARP1 at Asp214 within the DBD to generate fragments of 24 and 89 kD, creating a neo-epitope that is a hallmark of apoptosis.

To test whether PARP1 forms DNA adducts in response to treatment with 5-aza-dC, adducts were recovered and characterized from K562 cells, which derive from a chronic myelogenous leukemia patient in blast crisis, and from GM639 cells, which derive from SV40-transformed human skin fibroblasts [29]. These lines were chosen to represent myeloid tumors, which are treated with 5-aza-dC, and solid tumors, which are not. Small molecule inhibitors of PARP1 catalytic activity (PARPi) prevent PARylation, and several PARPi are currently FDA-approved treatments for some kinds of solid tumors [30, 31]. In some contexts 5-aza-dC synergizes with PARPi in cell killing [12, 13], so we also tested the response to PARPi, BMN-673 (known in the clinic as talazoparib), alone or together with 5-aza-dC. Control dose-response analyses showed that K562 cells were sensitive to 5-aza-dC and to PARPi, and no more sensitive to 5-aza-dC+PARPi combined than to either as a single agent; while GM639 cells were considerably more sensitive to the combined drugs (**Supplementary Fig. S1**). Western blots of whole cell extracts (**Supplementary Fig. S2**) established that 5-aza-dC treatment resulted in rapid degradation of DNMT1 and induction of the apoptotic 89 kD PARP1 fragment in both K562 and GM639 cells, as documented for other mammalian cell types [3].

Adduct recovery was carried out using a modification of the RADAR extraction method developed in our laboratory, in which cells or nuclei are treated with a combination of chaotropic salts and detergent to disrupt non-covalent protein-DNA interactions, and then nucleic acids, including covalent DNA-protein complexes, precipitated with alcohol [24-26]. K562 or GM639 cells were treated with 1 µM 5-aza-dC or 1 µM PARPi; or pretreated with for 48 hr with 5-aza-dC, allowing time for its incorporation into newly replicated DNA, after which PARPi was added and culture continued; followed by isolation of nuclei, RADAR extraction and precipitation of DNA and adducted proteins. Extracts were characterized by Western blots probed with antibodies tested for recognition of full-length PARP1 and its apoptotic fragments (**Fig. 1A**), to establish whether PARP1-DNA adducts had formed and to address the possibility that adducts had been cleaved by apoptotic caspases.

Probing adducts recovered from K562 cells with anti-recombinant PARP1 (anti-rec-PARP1, Enzo) identified species of 89 kD and smaller (55-65 kD) in samples treated with 5-aza-dC (66 hr) or treated with 5-aza-dC prior to treated with PARPi; but not in samples treated with PARPi alone (**Fig. 1B, left**). Reprobing this same blot with anti-N-ter-PARP1, which recognizes full-length PARP1 and both the 24 kD and 89 kD products of caspase cleavage, showed that samples treated with 5-aza-dC alone contained 113 kD PARP1-DNA adducts as well as larger PARP1 species that appear to be PARylated, as they were not evident in cells treated with PARPi, which inhibits PARylation (**Fig. 1B, right**). The anti-N-ter-PARP1 antibody also identified adducted 113 kD PARP1 and smaller species in extracts of cells treated with PARPi alone (18 hr).

Adducts recovered from GM639 fibroblasts treated with 5-aza-dC, PARPi, or 5-aza-dC+PARPi were analyzed by duplicate RADAR/Western blots run in parallel and probed with either anti-C-ter-PARP1, which recognizes both full-length and caspase-cleaved PARP1; or anti-cc-PARP1, which specifically recognizes the apoptotic N-terminal neo-epitope of caspase-cleaved PARP1. These two mAbs produced nearly identical staining patterns (**Fig. 1C**). In samples from cells treated with 5-aza-dC alone or 5-aza-dC+PARPi, both mAbs identified 89 kD cleaved PARP1 and smaller species (45-65 kD). Detection with anti-cc-PARP1 mAb, which specifically recognizes the apoptotic N-terminal neo-epitope, identifies the smaller species (45-65 kD) as probable proteolysis products of the adducted apoptotic 89 kD fragment. Comparably faint staining was evident in samples from untreated cells and cells treated with PARPi.

The results of RADAR/Western blots thus confirm the hypothesis that PARP1 forms covalent DNA adducts in cells treated with 5-aza-dC.

### 3.2 Mapping genomewide locations of covalent protein-DNA adducts by Adduct-Seq

If 5-aza-dC treatment causes PARP1-DNA adducts to form at hemi-methylated CpG dinucleotides that are targets of DNMT1 (**Fig. 2A, left**), then adducted DNA will be enriched in CGI, about half of which are associated with elevated DNA methylation [32-35]. CGI in human cells are defined by high GC fraction (>50%) and enrichment of CpG dinucleotides (http://genome.ucsc.edu/cgi-bin/hgTrackUi?g=cpgIslandExt) and by spatial position. These features are readily quantified by genomic sequencing.

In order to recover adducted genomic DNA fragments suitable for sequencing, we developed an approach, “Adduct-Seq”, that relies solely on endogenous covalent DNA-protein crosslinks to recover protein-bound DNA (**Fig. 2A, right**). RADAR extracts were prepared as for ChiP-Seq, but without addition of formaldehyde or other exogenous treatments to induce protein-DNA adduct formation. Samples were sonicated to reduce DNA length, then PARP1 and covalently bound PARP1-DNA adducts were captured by anti-C-ter-PARP1 mAb bound to a solid surface. Following deproteinization, recovered DNA was analyzed by genomic sequencing.

Samples were prepared from K562 or GM639 cells, untreated or treated with 5-aza-dC, PARPi, or 5-aza-dC+PARPi. From each sample of 1.5 x 10^7^ cells, 1.5-2.3 ng of PARP1-adducted DNA was recovered. Sequencing on an Illumina platform generated from 4.5-5.0 x 10^7^ mapped reads per sample (**Fig. 2B**). Samples from untreated cells were used to identify and assess background interactions and thereby distinguish the effects of drug treatment from intrinsic affinity of PARP1 for specific genomic regions. The untreated sample thus provides a control for background analogous to the “Input” sample in ChIP-Seq.

### 3.3 Adduct-Seq demonstrates PARP1-DNA adduct enrichment at CGI

Analysis of nucleotide composition of sequenced libraries (**Fig. 3A**) showed that the GC fraction of untreated samples was 42.0 (K562) and 42.5% (GM639), very close to the 41% average of the human genome. The GC fractions of adducts recovered from K562 cells treated with 5-aza-dC or 5-aza-dC+PARPi were significantly greater than that average (45.5% and 43.8% respectively; p<0.0001, Mann-Whitney test); while the GC fraction of adducts recovered from cells treated with PARPi alone was slightly lower than that average (39.7%). The GC fractions of adducts recovered from GM639 cells treated with 5-aza-dC or 5-aza-dC+PARPi were also significantly above the genomic average (47.2%, and 52.1%, p<0.0001, Mann-Whitney test), with the latter exceeding the 50% threshold defined for human CGI; while the GC fraction of adducts recovered from GM639 cells treated with PARPi alone (42.7%) was comparable to the genomic average.

**Fig. 3.**
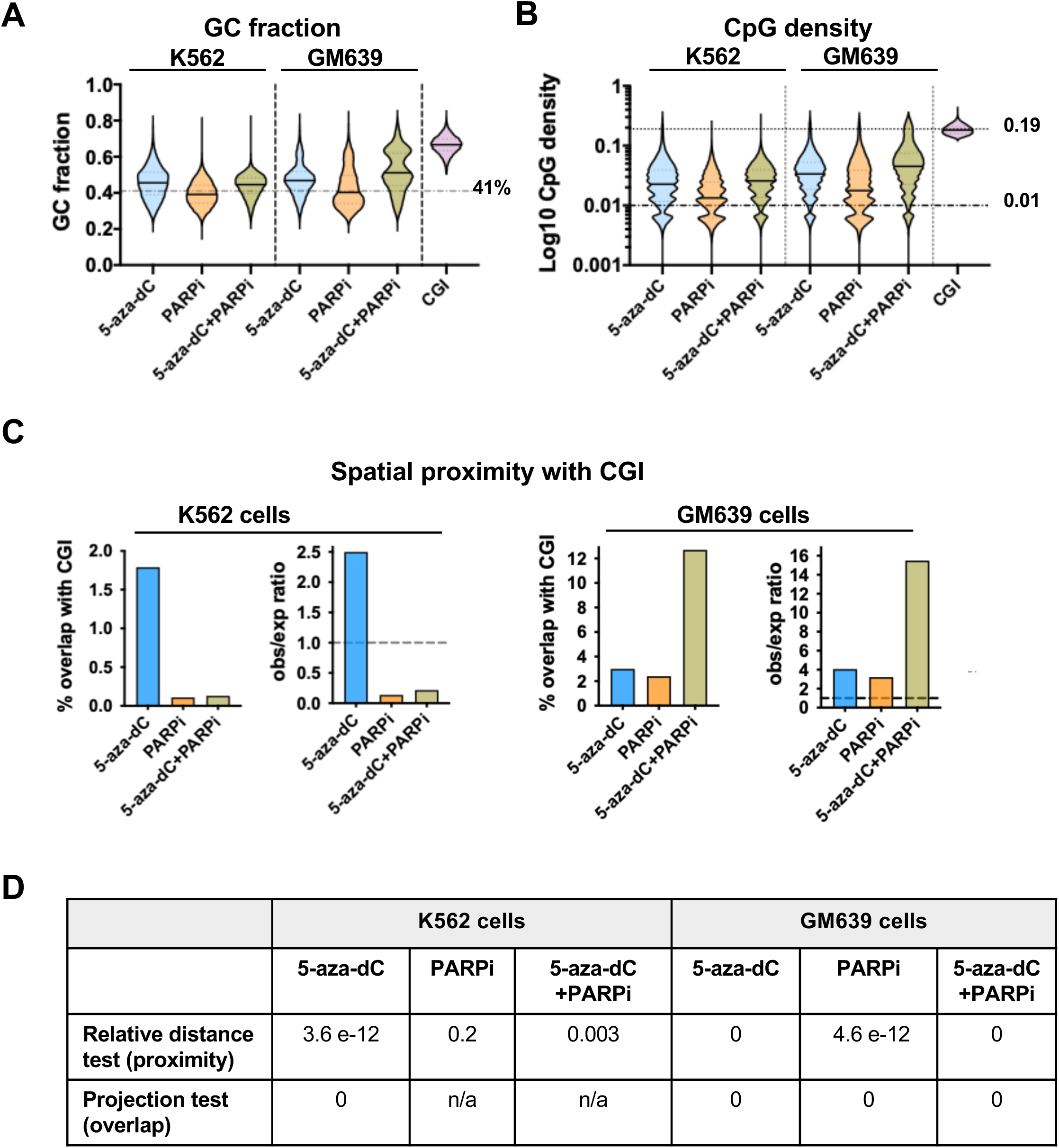
Genomic features of PARP1-DNA adducts recovered from drug-treated cells. (A) Violin plots of GC fraction of adducted DNAs recovered from K562 or GM639 cells after indicated treatments. Genome average GC fraction indicated by a dashed line. (B) Violin plots of CpG density of adducted DNAs recovered from K562 or GM639 cells after indicated treatments. Mean values of PARP1 peaks and CGI are marked by solid lines on violin plots; genomic average CpG density indicated by dashed line; mean CpG density of CGI indicated by a dotted line. (C) Results of spatial proximity analysis of DNAs analyzed by Adduct-Seq, showing % overlap with CGI in absolute values (left) and observed/expected (obs/exp) ratios (right) for each cell type. Dashed lines, ratio = 1. (D) Spatial correlation p-values for Adduct-Seq peaks and CGI locations shown in panel C. n/a – insufficient overlap for test to be applied.

Mean CpG densities of adducted DNA were determined (**Fig. 3B**) and compared to the human genome, where average CpG density is close to 0.01 and reaches 0.19 in CGIs. Mean CpG densities of adducted DNA recovered from K562 cells treated with 5-aza-dC or 5-aza-dC+PARPi were similar (0.030 and 0.028, respectively), significantly greater than either the genomic average or mean CpG density of adducts recovered from cells treated with PARPi alone (0.018; p<0.0001, two-tailed Mann-Whitney test). Mean CpG densities of adducted DNA recovered from GM639 cells treated with 5-aza-dC, PARPi or 5-aza-dC+PARPi were all significantly greater than the genomic average (0.041, 0.029, and 0.060 respectively; p<0.0001, two-tailed Mann-Whitney test).

Analysis of spatial correlation (**Fig. 3C-D**) of PARP1 peaks and the 31,144 CGIs in the human genome (CGI track in the UCSC Genome Browser) identified highly significant proximity and overlap among adducts recovered from K562 lymphoblasts treated with 5-aza-dC (relative distance test p=3.6e-12, projection test p=0). Significant proximity to CGI also characterized adducts recovered from K562 cells treated with 5-aza-dC+PARPi (p=0.003); but not adducts recovered from K562 cells treated with PARPi alone (p=0.2). Among adducts recovered from GM639 fibroblasts, highly significant proximity and overlap with CGI was observed in samples treated with either single agent 5-aza-dC or PARPi (relative distance test p=4.6e-12 and 0, projection test p=0 and 0, respectively), and combined treatment with both 5-aza-dC and PARPi further shifted adduct location towards CGI.

Adduct-Seq thus demonstrates that PARP1-adducted DNA fragments isolated from 5-aza-dC-treated K562 and GM639 cells exhibit significant enrichment for genomic features characteristic of targets of DNMT1 methylation: elevated GC fraction and CpG density and spatial overlap with CGI (**Fig. 3**). Genomewide analysis thus supports the hypothesis that PARP1 forms covalent adducts at sites of DNMT1 activity in 5-aza-dC-treated cells.

## 4. Discussion

The results reported here identify a plausible mechanistic pathway for cell killing by 5-aza-dC. **Fig. 4** outlines a simple working model for this pathway. In the course of normal replication, 5-aza-dC is incorporated opposite 5-me-CpG. DNMT1 is recruited to methylate the newly incorporated base, but the 5-aza group renders this a suicide substrate at which the adducted DNMT1-DNA intermediate persists, and its release damages the base. Repair generates an AP site or 5’-dRP that recruits PARP1, which forms an adduct that blocks replication. This activates apoptotic caspases 3/7, which cleave adducted PARP1.

**Fig. 4.**
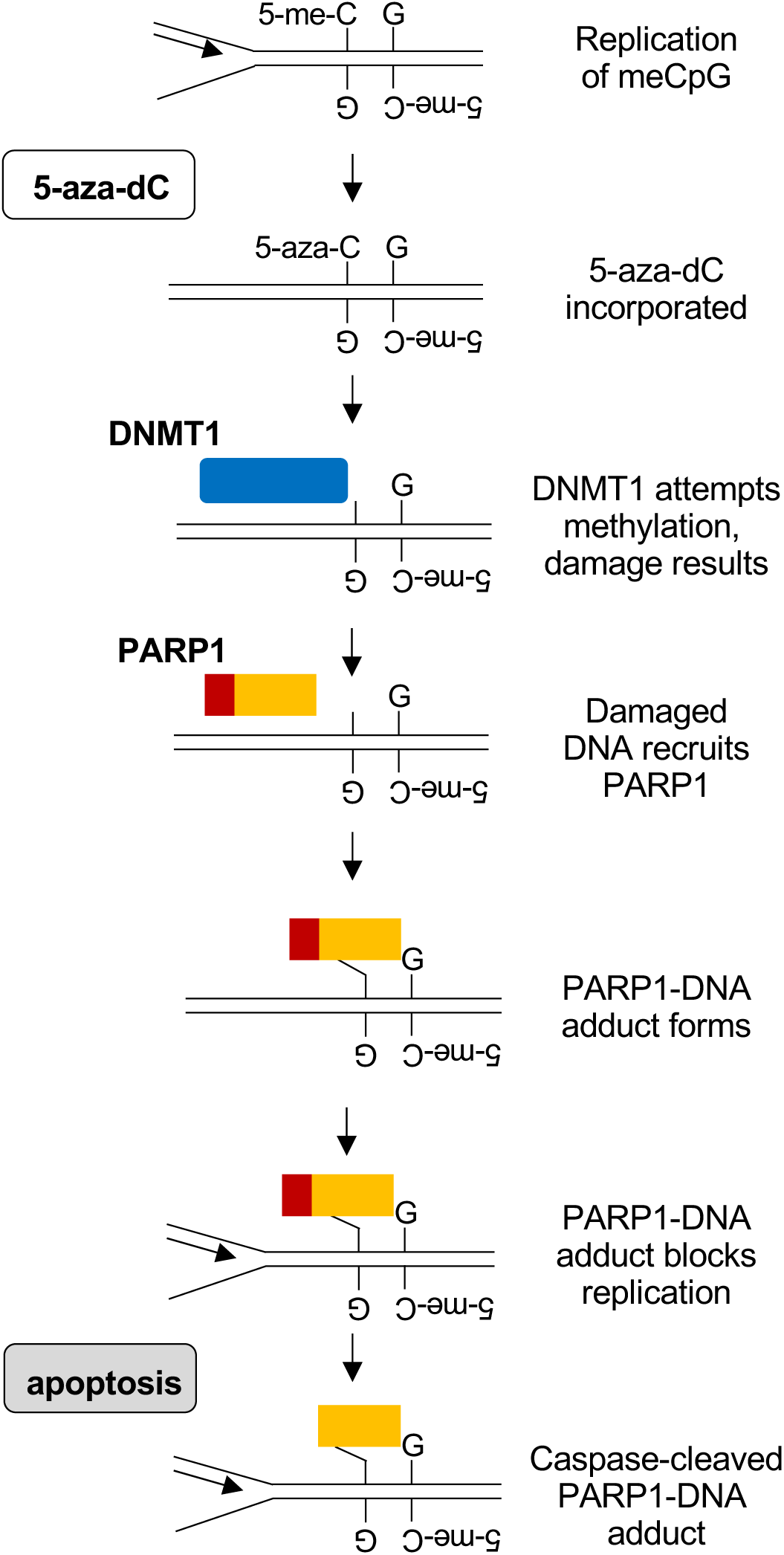
Working model for induction of cytotoxic PARP1-DNA adducts by 5-aza-dC. Proliferation in the presence of 5-aza-dC results in incorporation of 5-aza-C opposite a methylated CpG dinucleotide. Thwarted DNMT1 activity at this site results in DNA damage. PARP1 is recruited to the damage, where it forms a DNA adduct that — if not repaired — may cause replication arrest and cell killing, accompanied by activation of caspase 3/7 which cleaves adducted PARP1, creating the signature N-terminal neo-epitope.

The model specifically postulates that PARP1-DNA adducts form at the damage created by 5-aza-dC at hemi-methylated CpG dinucleotides that are targets for DNMT1. This was established by Adduct-Seq analysis (**Fig. 2**) of PARP1-adducted DNA fragments isolated from 5-aza-dC treated K562 and GM639 cells. Adducted fragments exhibit genomic features consistent with enrichment for targets of DNMT1 methylation: significantly elevated GC fraction and CpG density, and significant spatial overlap with CGI (**Fig. 3**). These results support a model in which aberrant activity of DNMT1 at 5-aza-substituted regions creates damage that recruits PARP1 to form covalent adducts, which arrest replication and induce apoptosis, activating caspase 3/7 cleavage of PARP1 (**Fig. 4**). Caspase cleavage generates the hallmark neo-epitope readily detected by RADAR-Western blots (**Fig. 1**), and caspase-cleaved PARP1 is disabled for critical activities including DNA damage recognition and DNA damage dependent PAR synthesis, and particularly chromatin recruitment [21-23, 36, 37]. While we cannot rule out the possibility that PARP1 was cleaved by apoptotic caspases prior to adduct formation, our results better support a model in which aberrant activity of DNMT1 at 5-aza-substituted regions creates damage that recruits PARP1 to form covalent adducts, which arrest replication and induce apoptosis.

The results reported here suggest the importance of revisiting the commonly held view of 5-aza-dC as an “epigenetic drug” [4, 5]. While treatment with 5-aza-dC undoubtedly affects genomewide methylation, induction of covalent adducts that block replication represents a straightforward path to cell killing that deserves further investigation. The results reported here further suggest that PARP1-DNA adducts may contribute to cell killing not only by 5-aza-dC, but also to other drugs, including PARPi, which can be readily tested by Adduct-Seq. If so, this may open the way to new therapeutic combinations.

## Acknowledgments

This research was supported by the US NIH National Cancer Institute (R01 CA183967 and P01 CA077852 to N.M.). We are grateful to Feinan Wu at the Fred Hutch, Seattle for assistance in analysis of Adduct-Seq results; and to Dr. Rajendra Prasad, National Institute of Environmental Health Sciences, for advising us on the antibodies for PARP1 adduct analyses.

## Supplementary Figure Legends

**Fig. S1.**
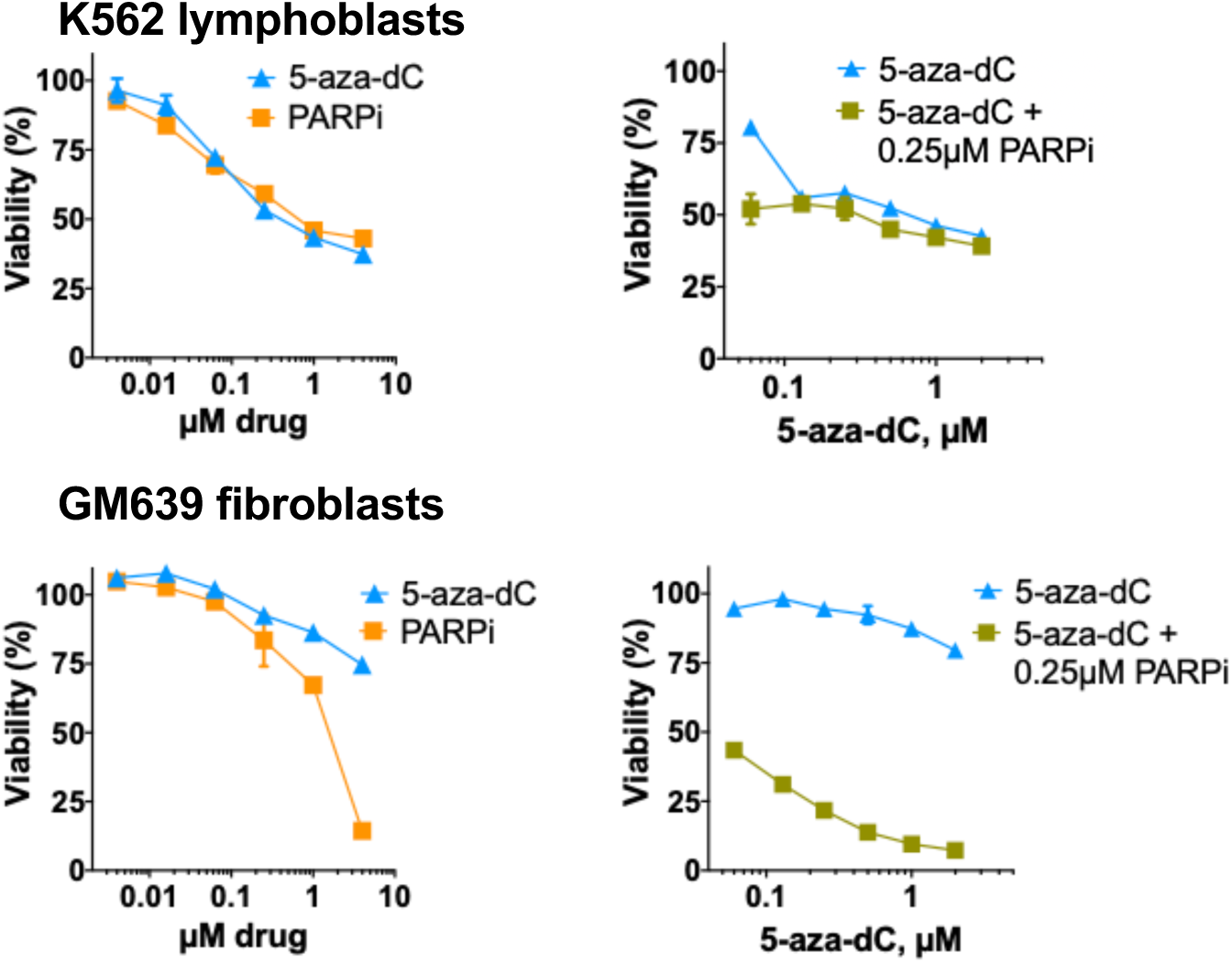
Responses of K562 and GM639 cells to drug treatment. Dose-response analysis of viability of K562 or GM639 cells treated for 96 hr with indicated concentrations of drug(s). Cells were incubated for 24 hr with 5-aza-dC prior to addition of PARPi. Viability assays were repeated twice, a representative experiment is shown.

**Fig. S2.**
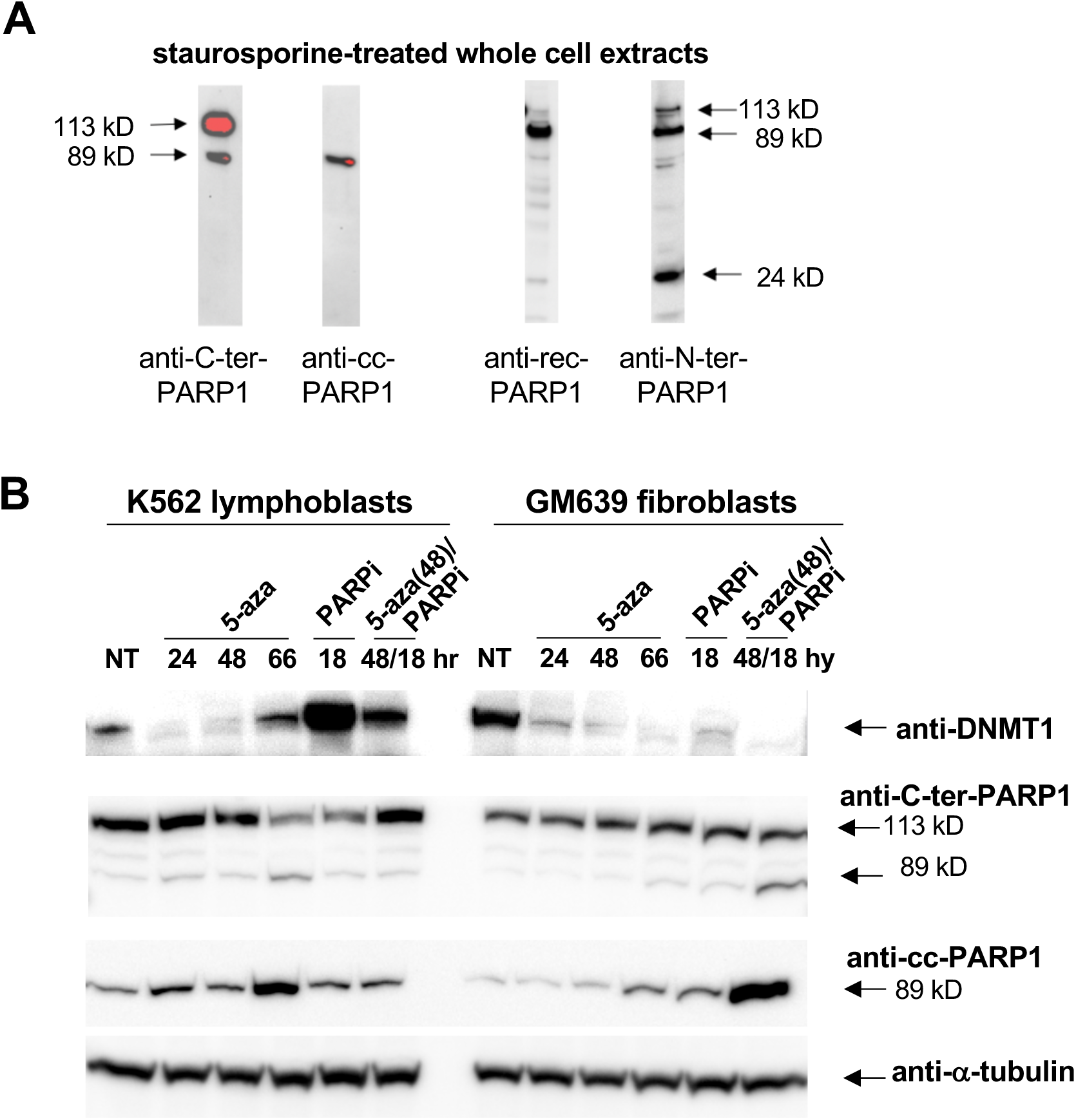
Responses of PARP1 and DNMT1 to drug treatment. (A) Anti-PARP1 Abs were tested for recognition of full-length 113 kD PARP1 and its cleavage products by Western blots of extracts from cells treated with staurosporine (STS; EMD Millipore; 1 µM, 1 hr). Blots were probed with anti-rec-PARP1, a rabbit polyclonal raised against the recombinant PARP1, 1:4000 dilution; anti-N-ter-PARP1, a rabbit polyclonal raised against the N-terminal half of recombinant PARP1, 1:2000 dilution; anti-C-ter-PARP1, a mouse mAb recognizing an epitope within C-terminal NAD binding/catalytic domain, 1:500 dilution, or anti-cc-PARP1, a mouse mAb specific for the neo-epitope at the N-terminus of the 89 kD PARP1 fragment formed after cleavage by apoptotic caspases between Asp214/Gly215, 1:1000 dilution). (B) Western blot analyses of extracts of K562 lymphoblasts and GM639 fibrobasts which were not treated (NT) or treated with 1 µM 5-aza-dC (5-aza; 24, 48 and 66 hr), 1 µM BMN-673 (PARPi, 18 hr), or pre-treated with 5-aza-dC for 48 hr prior to addition of PARPi and further culture for 18 hr (5-aza(48)/PARPi). The blot was probed with mAbs specific for DNMT1; the PARP1 C-terminal (anti-C-ter-PARP1); the neo-epitope on caspase-cleaved PARP1 (anti-cc-PARP1); or *α*-tubulin.

## Notes

### Competing Interest Statement

The authors have declared no competing interest.

